# Capturing context-specific regulation in molecular interaction networks

**DOI:** 10.1101/254730

**Authors:** Stephen T Rush, Dirk Repsilber

## Abstract

**Motivation:** Gene expression changes over time in response to perturbations. These changes are coordinated into functional modules *via* regulatory interactions. The genes within a functional module are expected to be differentially expressed in a manner coherent with their regulatory network. This perspective presents a promising approach to increase power to detect differential signals as well as for describing regulated modules from a mechanistic point of view.

**Results:** We present an effective procedure for identifying differentially activated subnetworks in molecular interaction networks. Differential gene expression coherent with the regulatory nature of the network is identified. Sequentially controlling error on genes and links results in more efficient inference. By focusing on local inference, our method is ignorant of the global topology, and as a result equally effective on exponential and scale-free networks. We apply our procedure both to systematically simulated data, comparing its performance to alternative methods, and to the transcription regulatory network in the context of particle-induced pulmonary inflammation, recapitulating and proposing additional candidates to some previously obtained results.

## 1 Introduction

Gene expression changes over time and in response to perturbation events, for example changes in environmental gradients. These changes are coordinated *via* regulatory interactions. In this article we identify differentially expressed subnetworks coherent with the regulatory structure, achieved by integrating differential gene expression with the associated network. Gene expression is routinely measured at the level of expressed RNA transcripts for each gene. Differentially expressed (DE) genes are those genes exhibiting a change in mean gene expression between conditions (see Figure 1). However, genes do not act in isolation. Rather, they act in biological networks consisting of interacting coordinated modules and more loosely coupled super-modules (Barabási and Oltvai, 2004). Ravasz *et al*. (2002) first demonstrated this empirically in organisms spanning the three domains of life, finding that their metabolic networks are organized into highly connected modules, which are then more loosely coupled in a hierarchical fashion. The genes within a functional module are expected to be differentially regulated in a coherent manner, i.e. respecting the regulatory network structure, in response to changes in their environment. From a systems level perspective, genes always act together in pathways and modules. The behaviour of these interactions aid in the study of the functions of genes and their products. For example, coordinated changes may be captured by gene co-expression patterns, which measure correlations. Use of direct correlations results in many false positives, and various methods exist to correct this (Friedman *et al*., 2008; Margolin *et al*., 2006). More recent methods profit from prior topological knowledge to constrain inference in network regulation. Specifically, there is an emphasis on context-specific network regulation (Warsow *et al*., 2010; Woo *et al*., 2015; Ernst *et al*., 2017; Hill *et al*., 2017). Numerous network-based regularization methods profiting from previous studies have emerged to perform variable selection and to obtain biologically meaningful predictors (Li and Li, 2008; Sun and Wang, 2012; Avey *et al*., 2017). Ma *et al*. (2016) perform gene enrichment analyses using either complete or incomplete topological information. These methods assume that a functional pathway is differentially active if most genes in this network structure are DE.

**Figure 1:**
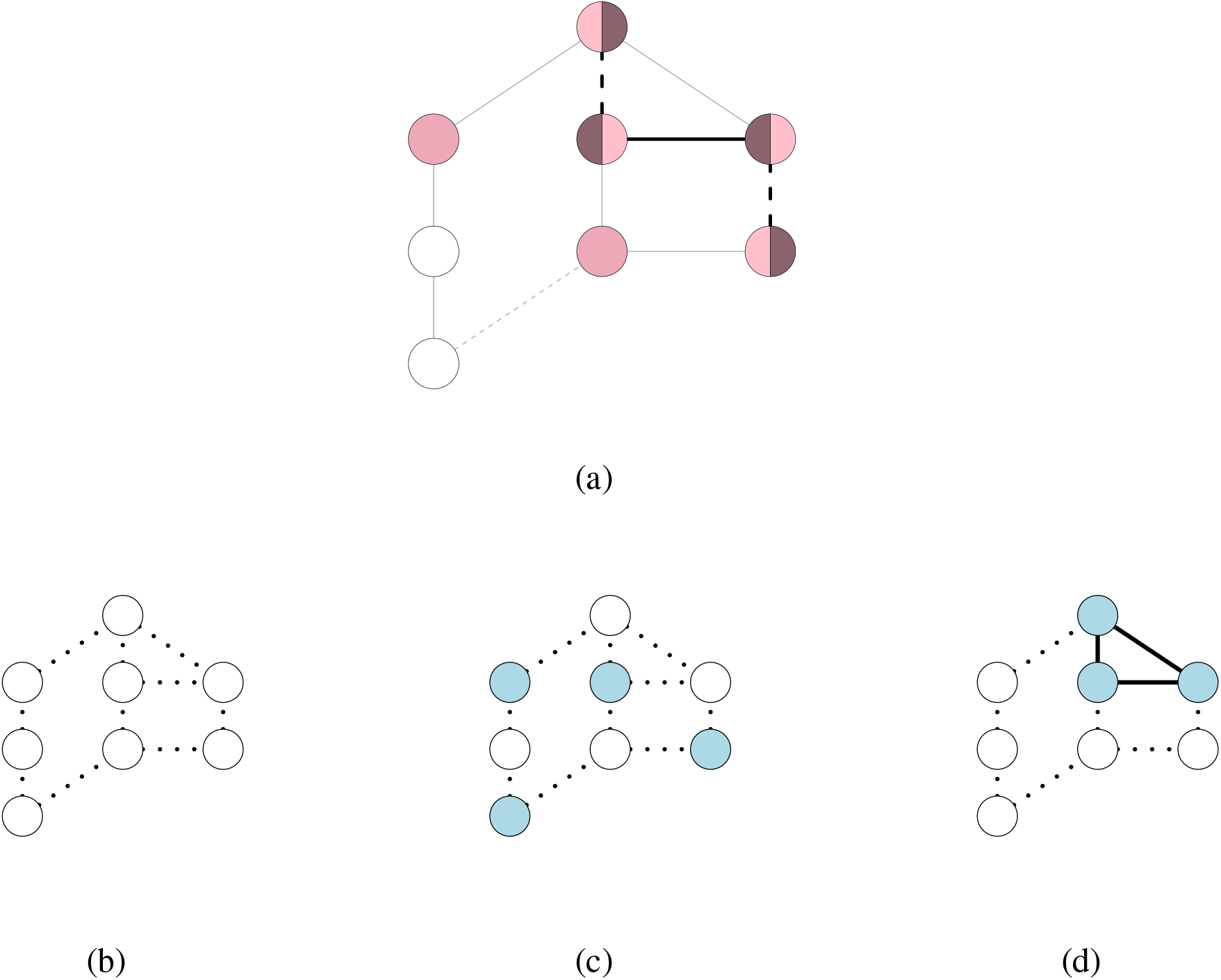
Differentially expressed (DE) genes. An example regulatory network. (a) The links between genes represent regulatory relationships: inducing (solid) and inhibitory (dashed). Some genes are DE, while others are not: DE-genes either increase (dark/light) or decrease (light/dark) in expression. Only some links are coherently expressed (thick edges). Note the non-DE boundary genes (solid). (b) Genes may not be DE. (c) Genes may be DE at random. (d) Genes may be DE in connected sub-modules.

In this article, we emphasize regulatory coherence. Regulatory coherence refers to gene expression patterns that respect the regulatory nature of the network. A network is described by a graph *G* = (*V, E*) for a set *V* of vertices and a set *E* of edges between vertices. For a gene regulatory network (GRN), the vertices represent genes while the edges indicate interactions between genes, such as activation, inhibition, modification, etc (see Figure 1a). We will refer to edges and vertices as links and genes, respectively. Inducing and inhibiting links are called regulatory links. Each gene regulates or is regulated by genes in its network topological neighbourhood. We define **coherent differential expression (CDE)** as the tandem changes in gene expression for a pair of genes in a link that is consistent with the regulatory nature of the link. We distinguish between inhibitory and non-inhibitory links. Non-inhibitory links consist of inducing links or relationships without explicit direction such as binding or positive correlations where the regulatory relationship is unknown. Tandem changes in gene expression for an inhibition link are coherent if, as the expression of gene *A* increases, the expression of gene *B* decreases. In contrast, CDE for non-inhibitory links occurs when the expression of both genes increases or decreases. Outside these two cases, differential expression is said to be incoherent. These concepts are illustrated in Figure 1a.

With regulatory coherence, it becomes clear that a GRN represents a collection of potential interactions, which are realized in specific contexts. We present a new perspective on the differential link score (Ernst *et al*., 2017) as a measure of regulatory coherence, along with a methodology to ascertain coherence of differentially active networks. These realized interactions form the **coherent subnetwork**. We systematically evaluate our methodology through simulation where the ground truth is known and through comparison with other methods for identifying differential expression in networks. Once validated, we apply our method to the problem of identifying DE subnetworks in an *in vitro* pulmonary inflammation study.

## 2 Methods

### 2.1 Coherent Differential Expression

In this section, we make our concept of coherence precise and develop an error control framework for identifying coherent subnetworks. Let *G* be a network with vertex set *V*(*G*) and edge set *E*(*G*), and let *S* ⊆ *G* be the coherent subnetwork corresponding to DE-genes, such as exhibited in Figure 1a. The genes in the vertex set *V*(*S*) are DE while all others, *V*(*G*)\*V*(*S*), are not. The edge set *E*(*S*) consists only of those links respecting regulatory coherence. It is not necessary that all edges between vertices in *V*(*S*) be included.

#### 2.1.1 Differential Link Score

Let *X_i_* ∈ ℝ^*p*^ be a vector of gene expression, *i* = 1,…,*n*, with number of genes *p*. Suppose there are two treatment groups, with labels 0 and 1. Without loss off generality, assume gene expression *X_i_, i* = 1,…,*n*_0_ < *n* corresponds to group 0 and gene expression *X_i_, i* = *n*_0_ + 1,…,*n* corresponds to group 1. Let *G* be the corresponding GRN with vertex *j* ∈ *V*(*G*) corresponding to gene expression *x_ij_*. For every gene pair (*j, k*) ∈ *E*(*G*), the differential link score (DLS) (Ernst *et al*. (2017)) is defined as

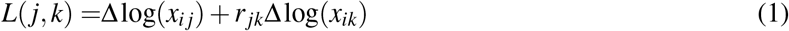

where *r_jk_* = –1 if gene *j* inhibits gene *k*, 1 otherwise. That is, the DLS is the sum of the log-fold change in expression of genes *j* and *k* adjusted for the sign of the regulatory relationship. Equation 1 can be re-arranged as the following:

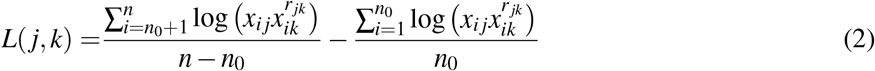

The terms 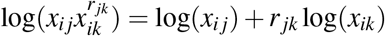 measure the simultaneous expression for two genes in a sample adjusted for regulatory relationship, which we call **coherent expression**.

Thus the DLS is the difference in mean coherent expression between classes. This captures regulatory coherence with respect to observed gene expression changes between the two classes. For inhibitory links, the DLS increases as both genes in the link change in opposition, whereas for non-inhibitory links, the DLS increases for simultaneous increases or decreases for both genes in the link.

#### 2.1.2 Hypothesis Testing

It is the second formulation of the DLS (Equation 2) which reveals the strategy for assessing significance of CDE for a gene pair (*j, k*) ∈ *E*(*G*). Observe that

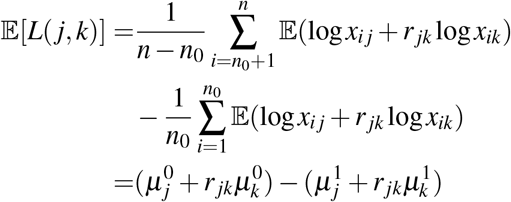

where 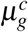 is the expected log-expression of gene *g* ∈ *V*(*G*) for class *c* = 0,1. To assess the significance of the link score *L*, we determine whether coherent expression is constant between classes. In other words, testing whether the mean coherent expression differs between two conditions is equivalent to testing whether their DLS is different from zero:

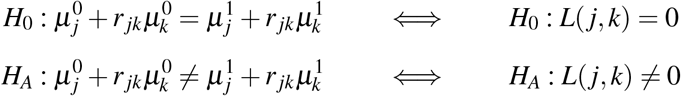

For two groups, independent samples, compute the coherent expression terms 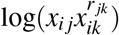 and then perform a two-sample test between classes. In this paper we will consider Student’s two-sample t (parametric) and Wilcoxon rank-sum (non-parametric) tests.

#### 2.1.3 Error Control

If we consider the definition of the DLS, we notice that *L*(*j, k*) = 0 implies either that neither gene is DE, or that both are DE to the same degree but in an incoherent way. However, if one gene is DE while the other is not, then *L*(*j, k*) ≠ 0. We call all links of this nature **boundary links** since they are found at the boundary of the differentially active module (see Figure 1a), and denote the collection of boundary links *∂S*. To discriminate between boundary links and links where both genes are DE, we need to identify both DE-genes and coherently expressed links.

We propose three methods of type I error control to identify DE-genes and CDE-links separately: (i) simultaneous control of genes and links (simultaneous), (ii) sequentially controlling genes and then links (forward), and (iii) sequentially controlling links and then genes (reverse). These are illustrated in Figure 2. We adopt the usual Benjamini and Hochberg (1995) false discovery rate (FDR) procedure for error control in all our analyses. A natural division of hypotheses is into edge and vertex families. We justify this division by Efron (2008), who demonstrated that FDR analysis is robust to separating hypotheses into mutually exclusive families and performing separate analyses at the same control level.

**Figure 2:**
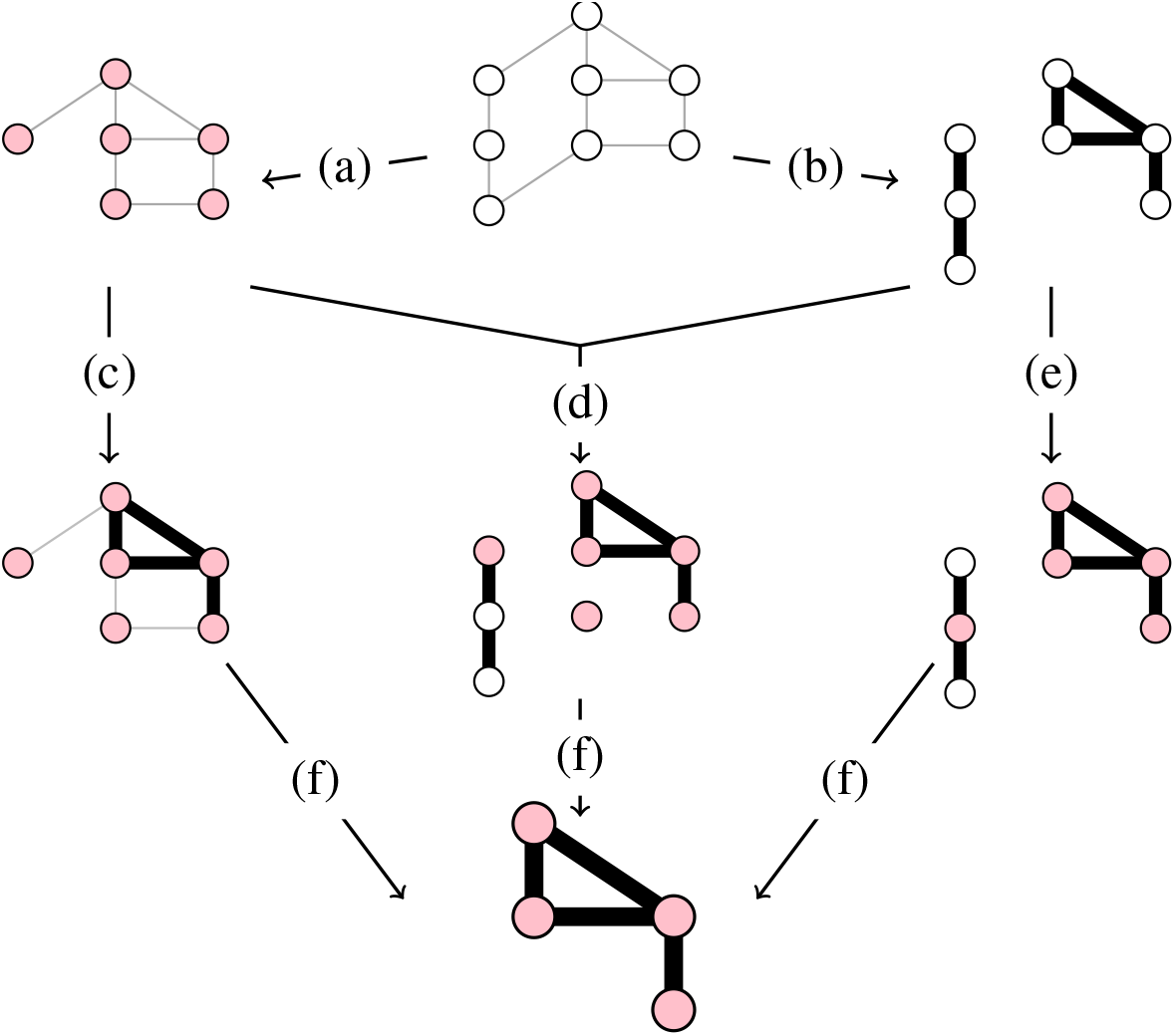
Coherent Differential Expression Procedures. The three error control procedures described in Section 2.1.3 are illustrated: Forward (acf), Simultaneous ([a+b]df), Reverse (bef). The steps are described as follows: (a) identify vertices; (b) identify edges; (c) identify edges, given vertices from a; (d) identify the subnetwork consistent with the edges and vertices selected at a and b; (e) identify vertices, given edges from b; (f) remove all isolated genes.

Let *F_V_* = *F_V_*(*G*) = {*H_v_*: *v* ∈ *E*(*G*)} be the family of tests concerning the vertices of the network *G*, and let *F_E_* = *F_E_*(*G*) = {*H_e_*: *e* ∈ *E* (*G*)} be the family of tests concerning the edges of the network *G*. Fix the nominal Type I error level at *α*. (i) For the simultaneous procedure, perform separate FDR analyses with *α*-level control for each family. Using the significant genes and links, identify the *α*-significant coherent subnetwork *S* ⊆ *G* and the boundary links *∂ S* separating the coherent subnetwork from the rest. (ii) For the forward procedure, perform FDR analysis on the vertex family *F_V_* first, then delete all non-significant vertices from the network *G*, obtaining the subgraph *S* ⊆ *G*. Then, perform a separate FDR analysis on the edge sub-family *F_E_*(*S*) ⊆ *F_E_*, in this way identifying the *α*-significant differential subnetwork. (iii) For the reverse procedure, perform FDR analysis on the edge family *F_E_* first, then prune all non-significant edges from the network *G*, obtaining the subgraph *S* ⊆ *G*. Then, perform a separate FDR analysis on the vertex sub-family *F_V_*(*S*) ⊆ *F_V_*, in this way identifying the *α*-significant coherent subnetwork. In all three procedures, we delete isolated genes.

### 2.2 Network Models

#### 2.2.1 Network Topology

We consider two classes of random graphs, the (first) Erdős-Rényi model and the Barabási-Albert model (Albert and Barabási, 2002). The Erdős-Rényi model considers an initial set of *v* vertices, with *e* edges chosen uniformly from the set of all *v*(*v* – 1)/2 unique edges between vertices. Its topology is said to be exponential due the distribution of vertex degree, which follows a Poisson distribution. On the other hand, the Barabási-Albert model belongs to the class of scale-free graphs, so-called because there is no ‘typical’ node degree, with the degree distribution following an approximate power-law. Beginning with the biologically compelling assumption that as a network grows, new nodes attach preferentially to nodes with higher degree, Albert and Barabási (2002) demonstrated that random graphs produced in this way are scale-free. Even though most biological networks appear to be scale-free, exponential graphs still arise naturally. Barabási and Oltvai (2004) mention for instance that *Saccharomyces cerevisae* and *Escherichia coli* exhibit mixed exponential and scale-free features, noting that the incoming degree distribution for transcription regulatory networks is approximately exponential while the degree distribution of transcription factor interactions is scale-free.

#### 2.2.2 Graphical Models

In a regulatory network *G*, each node *j* ∈ *V*(*G*) interacts with a subset of the network, its neighbourhood *N_j_* = {*k*: (*j, k*) ∈ *E*(*G*)}. Let *X_j_* be the random expression for a gene *j*. The expression *X_j_* is anticipated by its neighbourhood: given the expression in neighbourhood *N_j_*, no further information is gained for the prediction of *X_j_* by learning the expression of gene *l* ∉ *N_j_, l* ≠ *j*. In other words, nodes *j* and *l* are conditionally independent given *N_j_*. The joint distribution of gene expression may be factorized along the maximal cliques of the graph, and hence motivates the application of graphical models. We simulate differential gene expression using Gaussian graphical models (GGMs), which may be specified by their mean vectors and inverse covariance matrices (Kolaczyk and Csárdi (2014)). Let Σ^−1^ be an *n × n* real positive definite matrix such that (Σ^−1^)_*jk*_ = 0 whenever (*j, k*) ∉ *E* (*G*) for *j* ≠ *k*. This implies that the partial correlation between vertices *j* and *k* is zero for all non-linked vertices in the network. Further let μ be fixed in ℝ^*n*^. Then the distribution 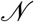(μ, Σ) describes a GGM with corresponding network *G*.

### 2.3 Simulation of Differential Expression in Networks

We develop simulations to evaluate the ability of our method to identify coherent interactions and DE modules. To model the dependence structure among genes, gene expression data is log-normally distributed according to a GGM. For each replicate, a random network *G*, covariance matrix Σ consistent with *G*, and mean log-expression vectors *μ*_0_,*μ*_1_ are generated. Log-expression is sampled from models *N*(*μ*_*c*_,Σ), *c* = 0,1. Each simulation consists of 100 replicates. We provide details in the following.

#### 2.3.1 Random Graphs

In the simulations we use the following random networks: (i) Exponential Erdős-Rényi and (ii) Scale-free Barabási-Albert. (i) For the exponential graphs, we supply the following parameters for the number of vertices and edges (*v,e*): (500,2000), and (2000,8000). (ii) For the scale-free graphs, we set the number of vertices, power of preferential attachment, and number of edges to add at each time-step (*v, p, m*) as (500,1,2) and (2000,1,2). See `igraph` for details (Csardi and Nepusz, 2006).

#### 2.3.2 Differential Expression and Localization Patterns

We investigate both null and true differential expression. For the small graphs (*v* = 500), we simulate data where (i) there is no differential expression (null), (ii) differential expression is distributed randomly over the vertices (low: 1% DE; high: 10% DE), (iii) differential expression is restricted to a connected subgraph (low: 1% DE; high: 10% DE), (iv) differential expression is restricted to three connected subgraphs (low: 3% DE; high: 30% DE) with average size 5 (low) and 50 (high). For the large graphs (*v* = 2000), we simulate data where (i) there is no differential expression (null), (ii) differential expression is distributed randomly over the vertices (low: 1% DE; high: 10% DE), (iii) differential expression is restricted to a connected subgraph (low: 1% DE; high: 10% DE), (iv) differential expression is restricted to three connected subgraphs (low: 1% DE; high: 10% DE) with average size 20 (low) and 200 (high). For each gene, mean log-fold differential expression is pulled from a uniform distribution *U*(–*d,d*), *d* = 4,8,16,32, corresponding to mean log-fold differential expression 2.7,5.3,10.7,21.3.

Expression patterns are depicted in Figures 1bcd. To obtain a connected subnetwork for each module, we perform the following procedure:

1. Select vertex *v*_1_ randomly from the vertex set of the graph *G*, and put this in *V* = {*v*_1_}.
2. Choose a vertex from the vertex neighbourhood of *V* in *G* and add it to *V*.
3. Repeat until obtaining the vertex set 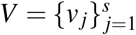, with *s* the desired size of the subset.

In the case of multiple modules, each module is created to be approximately the same size.

#### 2.3.3 Covariance Structure

The covariance structure must be informed by the graph structure of the network, as well as the nature of the link. In Eukaryotes, inducing links account for approximately 75% to 80% of regulators (McDonald *et al*., 2008; Wang *et al*., 2017). McDonald *et al*. (2008) report that the average proportion of activations for circadian networks is 0.74 in *Arabidopsis* and *Drosophila*, while generally for Eukaryotic signalling networks the average is 0.83. For each graph, we choose a random proportion uniformly over (0.72,0.85), *p* ∼ *U*(0.72,0.85). We assign each edge a relationship *r_ij_* of 1 (non-inhibitory) with probability *p* and −1 (inhibitory) otherwise.

We construct a covariance matrix satisfying the conditional dependence structure of the network by first constructing the precision matrix **P** = [**p_ij_**] from the adjacency matrix **A** = [**a_ij_**] and then inverting to obtain the covariance matrix Σ = **P**^−1^, described in Algorithm 2.1.

**Algorithm 2.1:**
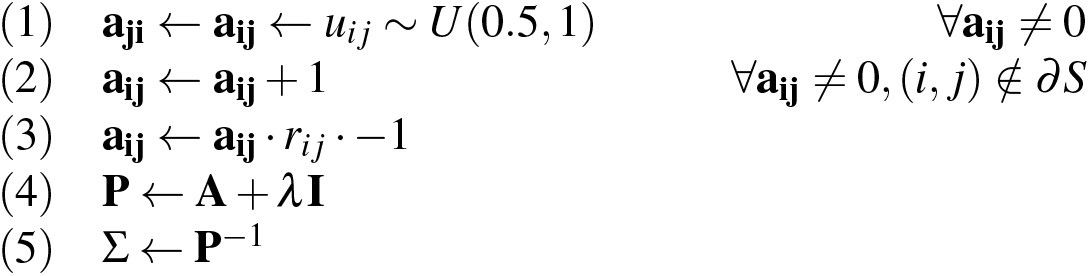
Generate Covariance Matrix(*R*)

In step (1), we assign random values to edge weights independently and identically distributed according to a uniform distribution *U*(0.5, 1). In step (2), we add 1 to all non-boundary edge weights, ensuring that the genes in modules are more strongly coupled to each other than to the rest of the network. In step (3), we adjust all edge weights by their relationship encoded in *r_ij_*, and multiply this by −1 to account for the relationship between partial correlation *ρ_ij_* and the entries of the precision matrix 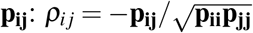. In step (4) we add a positive shift λ**I** to ensure positive definiteness. The scale parameter λ = λ_1_ + λ_2_ is calculated as follows. –λ_1_ is the smallest eigenvalue of the matrix **A** from step (3). It ensures positive semi-definiteness. We then calculate λ_2_ > 0 so that the resulting matrix has condition number equal to the number of vertices *v* in the network. This ensures invertibility of the matrix. Finally in step (5) we obtain the covariance matrix. In our simulations, we use the corresponding correlation matrix.

#### 2.3.4 Evaluation

Performance in simulations is evaluated via precision (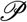), sensitivity (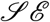), and specificity (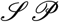) of the procedures to both genes and links. Explicitly, let *G* be the simulated regulatory network, *S* be the simulated coherent subnetwork, and *Ŝ* the estimated coherent subnetwork. Then

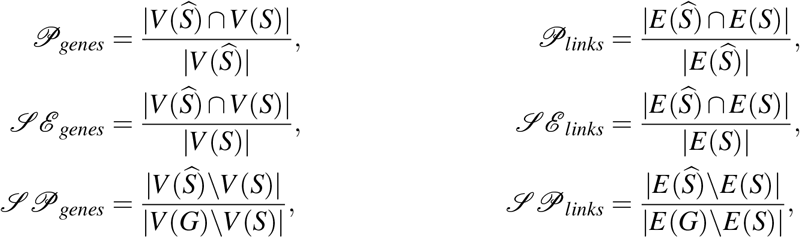

where | · | counts the number of elements in a set.

### 2.4 Alternative Methods

We compare the CDE procedures to three alternatives, a baseline network independent method and two methods incorporating network constraints.

Limma is a linear model based method that uses moderated t-statistics to assess the significance of the design as a predictor of gene expression (Ritchie *et al*. (2015)). In our simulation study, we use limma as the baseline network-free method. We ascertain the significance of gene expression log-contrasts 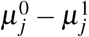 with Benjamini-Hochberg FDR (*α* = 0.05).

BioNet incorporates a network in the analysis of gene expression profiles for the detection of functional modules (Beisser *et al*. (2010)). Beginning with a set of *p*-values assigned to each vertex, a beta-uniform mixture model is fit. Scores for each subnetwork are computed based on this model and an integer linear programming algorithm is used to locate the maximum scoring subnetwork. For our simulation study, we take the unadjusted *p*-values obtained for limma and feed them into the BioNet algorithm with Benjamini-Hochberg FDR control (*α* = 0.05).

FocusHeuristics is a quantile-based method to highlight the differentially active components of a gene expression network in order to focus on phenotypically-relevant subnetworks (Ernst *et al*., 2017). The authors define three scores on the edges and vertices of the network: (i) log-fold change Δlog(*x_ij_*), (ii) differential link score *L*(*j,k*), and (iii) interaction link score min{log(*x*_0*j*_ + *x*_0*k*_),log(*x*_1*j*_ + *x*_1*k*_)}. All edges and vertices passing the quantile threshold for at least one of these three scores are retained. One difficulty with FocusHeuristics is the number of parameters to tune, for which no systematic decision rule is indicated. In order to compare our error-controlled procedures with FocusHeuristics, we restrict to a single replicate, evaluating the performance of the seven expression patterns. For FocusHeuristics, we choose the same thresholds for each of the scenarios by scaling the default thresholds so that the average size of the subnetwork is approximately 200 vertices.

### 2.5 Application

We consider a gene expression experiment investigating particle-induced inflammation in pulmonary artery endothelial cells reported in Karoly *et al*. (2007), specifically the effect of exposure to airborne ultrafine particles (UFPs) ‒ particles with diameter less than 100 nm. The authors hypothesize that UFPs contribute to endothelial cell dysfunction by inducing transcriptional activation of genes involved in coagulation and inflammatory responses. To test this, they perform a cell culture study with one treatment group (exposure to 100*μ*g/mL UFPs; n=4) and one control (no UFP exposure; n=4) and measure the effects via gene expression.

**Gene expression**. Affymetrix microarray `CEL` files are downloaded from the Gene Expression Omnibus (GEO) database (Barrett *et al*. (2013)), accession number GSE4567. Gene expression is corrected and normalized via the R-package `oligo` (Carvalho and Irizarry (2010)) using the default method and then log-transformed. Expression data is annotated with gene symbols using the R-package `hgu133plus2.db` (Carlson (2016)). Where gene symbols correspond to multiple expression values, we take the mean of the values within each sample.

**Gene regulatory network**. The human TRRUST V2 network (Han *et al*. (2018)) is used as the seed network. This network consists of 800 transcription factors (TFs) and 2,095 non-TFs, with 8,444 regulatory links. We remove loops and multiple edges. After restricting the gene expression dataset and TRRUST network to their common gene set, we obtain a GRN of 2,731 nodes and 7,966 links.

**Coherent differential expression**. We infer the coherent subnetwork via the parametric forward control (F/P) procedure using Welch’s t-test with Benjamini-Hochberg control (*α* = 0.05). We interpret the results using STRING (Szklarczyk *et al*., 2017), a database for protein-protein association networks. We compare our gene list to the *Homo sapiens* database and identify whether coagulation or inflammation pathways are enriched. STRING identifies enriched gene sets using Fisher’s exact test (Rivals *et al*. (2007)) with Benjamini-Hochberg FDR control.

## 3 Results and Discussion

### 3.1 Simulations

We present a subset of the simulations in the text. The full set of simulations are presented in the Supplement. The comparisons between alternative CDE-procedures are presented in Suppl. Figures 1-24, the benchmark comparisons are presented in Suppl. Figures 25-30, and the comparisons with FocusHeuristics are presented in Suppl. Figures 31-34.

**Error control on genes and links is independent of network topology**. We observe no differences in performance between the two network topologies, nor across network sizes. This suggests that these procedures are network-invariant, and therefore appropriate for more complex network structures.

**The false discovery rate is asymptotically controlled**. The empirical FDR of the simultaneous (S) and forward (F) procedures generally fall well below the nominal control level, while FDR for the reverse (R) procedure is controlled asymptotically as sample size or average change in expression increases when differential expression presents in modules; this pertains to both genes and links. Across all procedures, variation in FDR decreases as network size increases. This is explained by the increasing number of DE-genes. Indeed, we observe the same decrease in FDR variation as the DE proportion increases for fixed network size. FDR is poorly controlled by procedure R when DE is scattered randomly across the network. This is to be expected, since procedure R begins with identifying links, of which there should be few, and we delete all isolated genes. Therefore the cost in improved performance for differentially active modules is borne by a decrease in performance for random differential activation. The results for the full set of simulations are presented in Suppl. Figures 1-24.

**Measuring links first increases sensitivity at the cost of some loss in precision**. In general, procedure R is most sensitive while procedure S is least. However, as sample size and average differential expression increases, we sometimes observe that procedure F is more sensitive than procedure R. All three procedures converge on sensitivity as sample size and average differential expression increases. Sensitivity with respect to genes is poor for scattered differential expression. We expect this, since we deliberately exclude isolated genes.

**The parametric sequential control procedures outperform the rest**. In general, the parametric (P) procedures outperform the non-parametric (NP), although they are generally close. While specificity for P is typically less than NP, this is compensated with a higher sensitivity and a more faithful error control. This is expected, given that the t-test is more efficient when data are normal. Procedure F consistently outperforms procedure S. Procedure R typically dominates in sensitivity while procedure F dominates in error control. We consider only the parametric sequential control procedures forward parametric (F/P) and reverse/parametric (R/P) in the sequel.

**The procedures effectively identify modules**. There are almost zero false positives in the null differential expression scenario. This suggests that if several modules were DE, we could identify them and discriminate between them. Indeed, results for scattered and modular differential expression confirm this. Whereas we do not typically obtain large connected components for scattered differential expression, but only a few small components of two or three genes, we identify large modules in the modular differential expression scenario. Thus DE-modules identified by our procedures constitute actual findings.

**Error control on genes effectively removes the boundary**. One might expect boundary links to be over-represented among false positives, since by definition their link has nonzero coherent expression. However, we find that there is no great difference in specificity between scattered and modular differential expression for links. This is significant because the scattered differential expression case has a different boundary issuing from sparse connectivity of DE-genes. Thus controlling error on the vertices is sufficient to control these errors.

**The CDE procedures outperform the network-independent baseline and BioNet when differential expression is present in modules**. The performance of the limma baseline procedure is nearly independent of the differential expression pattern, with strict error control over all DE proportions, average change in differential expression, sample size, network size, and network topologies, with sensitivity increasing as sample size or average change in differential expression increases, see Figure 3. The CDE procedures are more sensitive when differential expression is modular, at the expense of performing poorly when differential expression is scattered. BioNet itself is outperformed by limma and the CDE procedures except for low sample sizes or low change in average differential expression. BioNet tends to control error similarly to the R/P procedure, with less variation in FDR; however its variation in sensitivity and to some extent specificity is more extreme.

**Figure 3:**
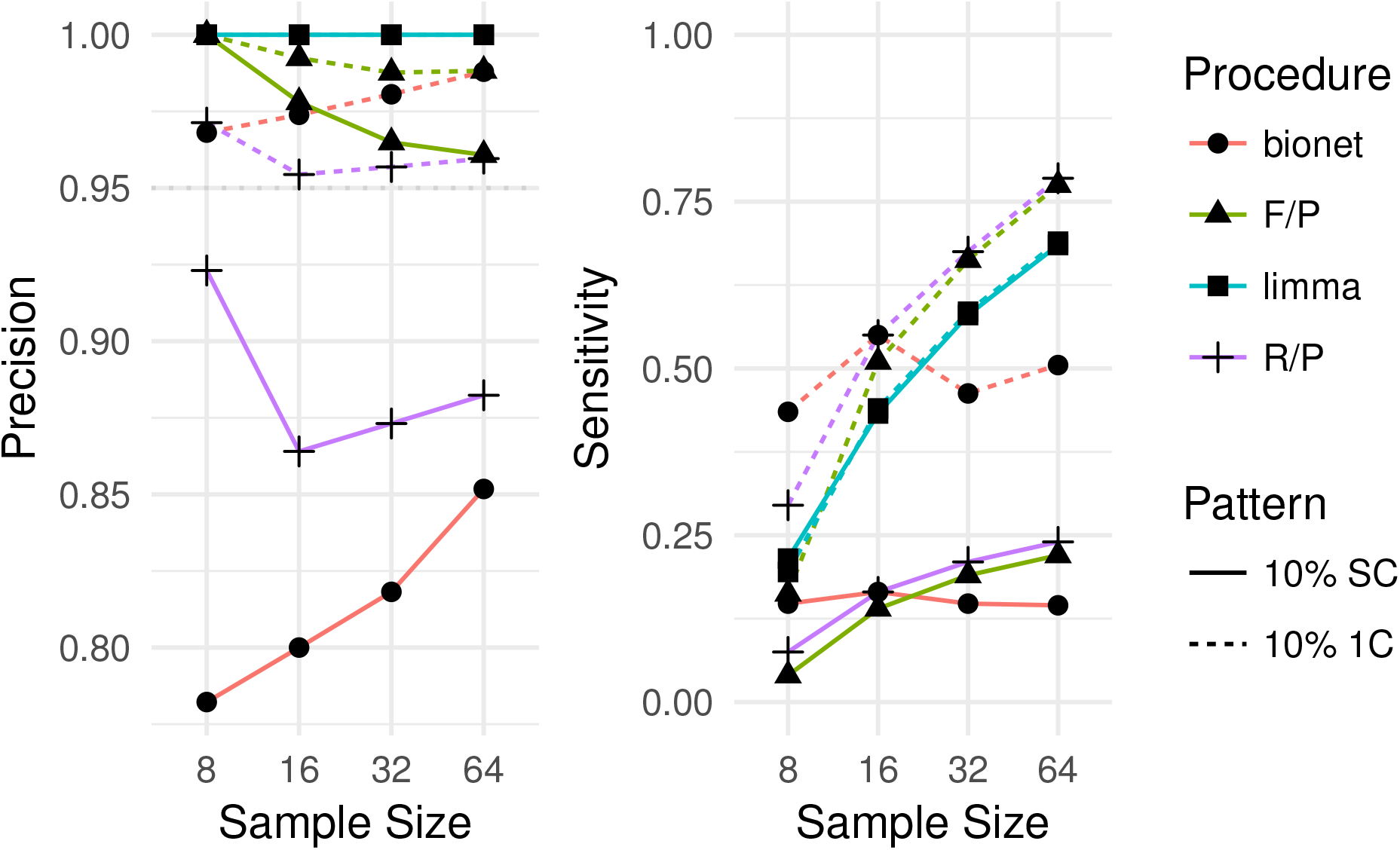
Performance of Differential Expression Procedures. We report the median results for large scale-free networks (100 replicates) with 10% differential expression. Displayed are the results for the Forward Parametric (F/P) and Reverse Parametric (R/P), BioNet, and limma procedures on the scattered (SC) and 1 module (1C) patterns.

**The CDE procedures outperform FocusHeuristics**. We immediately note that for our procedure the number of selected vertices increases as the active proportion increases (Figure 4a-c), whereas the number of vertices chosen by FocusHeuristics is independent of the active proportion (Figure 4d-f). For the scattered differential expression scenario, there should not be many adjacent DE genes. Despite this, FocusHeuristics results for the scattered differential expression scenario are indistinguishable from those for the modular scenario. In contrast, our error-controlled procedures identify more sparsely connected subnetworks for the scattered differential expression scenario and select no false positives in the noise scenario. While FocusHeuristics is able to identify some of the DE component of a system, it is unable to distinguish between signal and noise.

**Figure 4:**
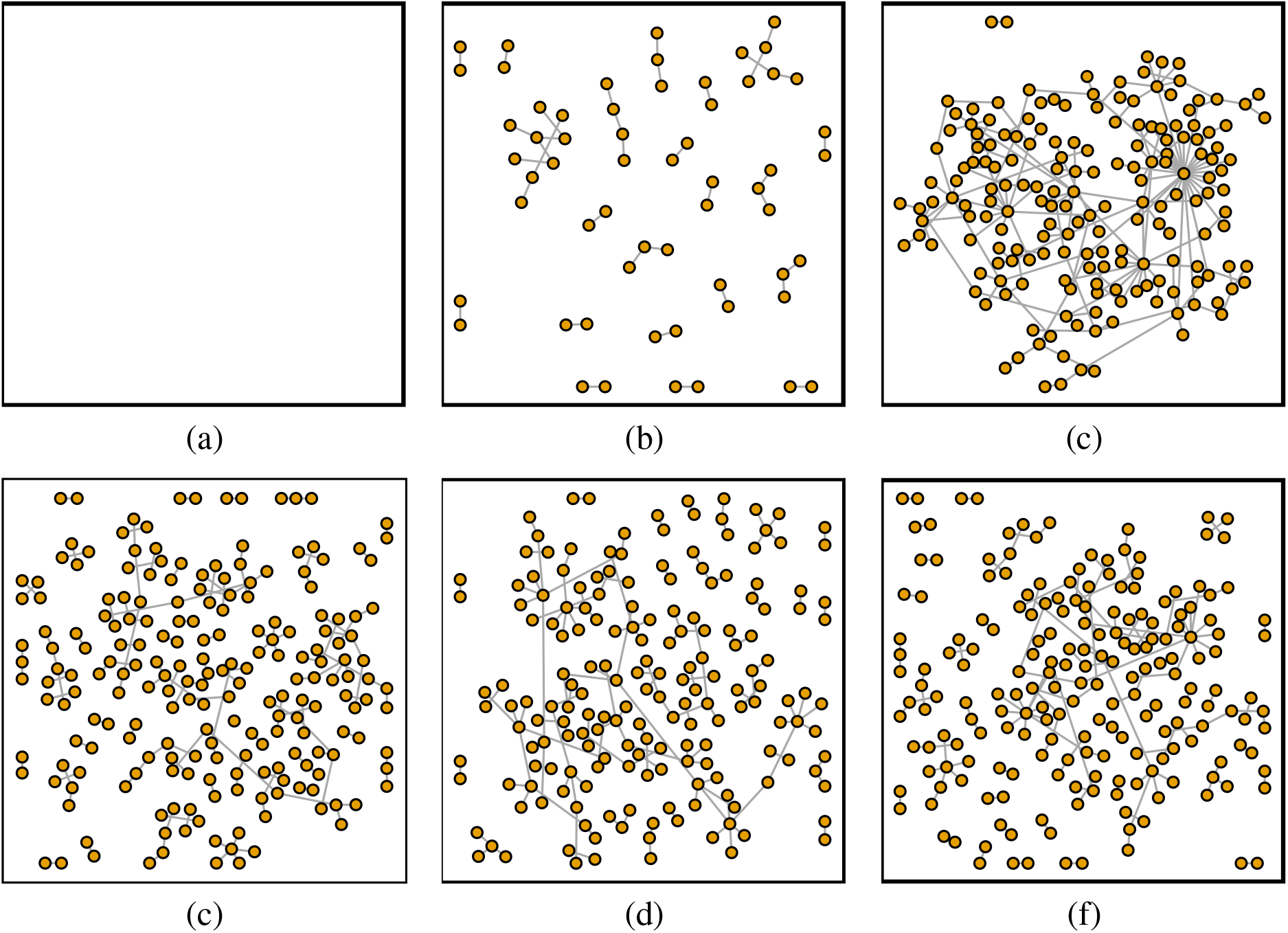
Controlled Coherent Differential Expression versus FocusHeuristics. Figures (a)-(c) illustrate the performance of the Forward Parametric (F/P) procedure on the 0% differential expression (DE), 10% scattered DE, and 10% single module DE scenarios, respectively. Figures (d)-(f) illustrate the performance of FocusHeuristics on the 0% differential expression, 10% scattered DE, and 10% single module DE scenarios, respectively. Notice that Figure (a) depicts an empty network; in this instance, the F/P procedure has identified no false positives.

### 3.2 Application: Particle-induced inflammation

We obtain a coherent subnetwork with 64 genes and 85 links, with one large connected component and 4 components consisting of gene pairs. we note that most genes in the coherent subnetwork increase in gene expression, indicating that the dominant response is activation. Thus we can speak of pathways being activated by exposure to UFPs. STRING identified 482 GO Biological Processes and 41 KEGG Pathways as significantly enriched (FDR *α* = 0.05), compared to Karoly *et al*. (2007) which identify three KEGG Pathways. While we identify the cytokine-cytokine receptor interaction KEGG pathway observed in Karoly *et al*. (2007), we fail to detect the Wnt signalling or MAPK signalling KEGG pathways, instead detecting other enriched pathways, a number of these corresponding to inflammation processes. We screened out all infection and cancer pathways, since these also lead to inflammation but are not relevant to the present study. This left 19 remaining pathways, 9 of which correspond directly with inflammation, 2 indirectly via oxidative stress, and 3 corresponding to other immune responses, see Table 1. The identification of oxidative stress pathways is consistent with the hypothesis that air-borne particles induce inflammatory response through an oxidative stress mechanism. Karoly *et al*. (2007) are particularly interested in tissue factor (TF), noting that its gene expression (F3) is up-regulated. In an additional experiment, they find that increased TF protein induces increased release of the cytokine IL-8. We can form the hypothesis that the gene F3 induces cytokine release (e.g. genes *CXCL*1, *CXCL*2, *IL*6, *CXCL*8) through its neighbours *EGR*1 and *NFKBI*, see Figure 5.

**Table 1:**
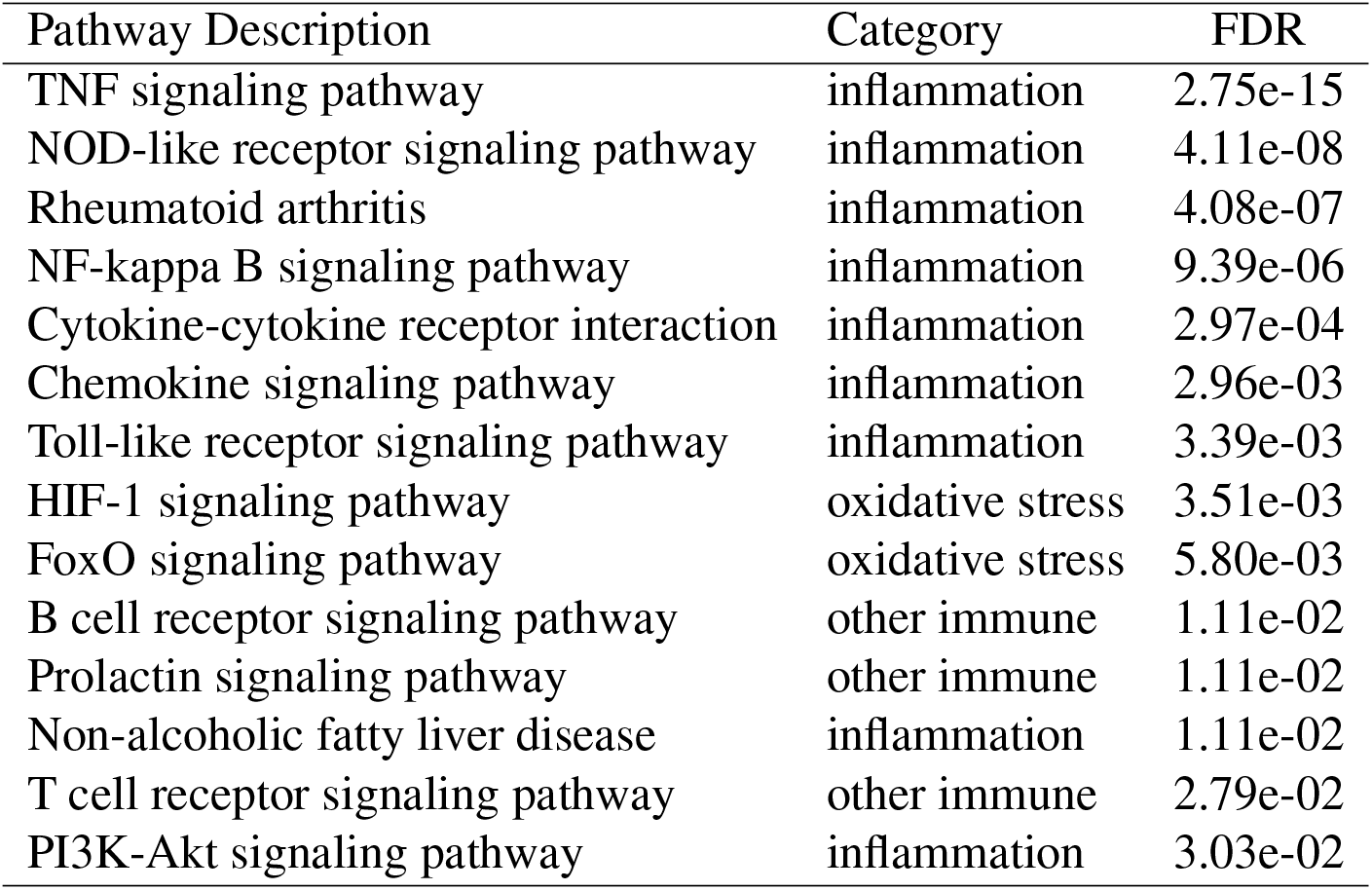
**Ultrafine particle-exposure coherent enriched pathways**. The non-infection, non-cancer, immunerelated KEGG pathways identified as enriched by STRING using Benjamini-Hochberg adjusted p-values.

**Figure 5:**
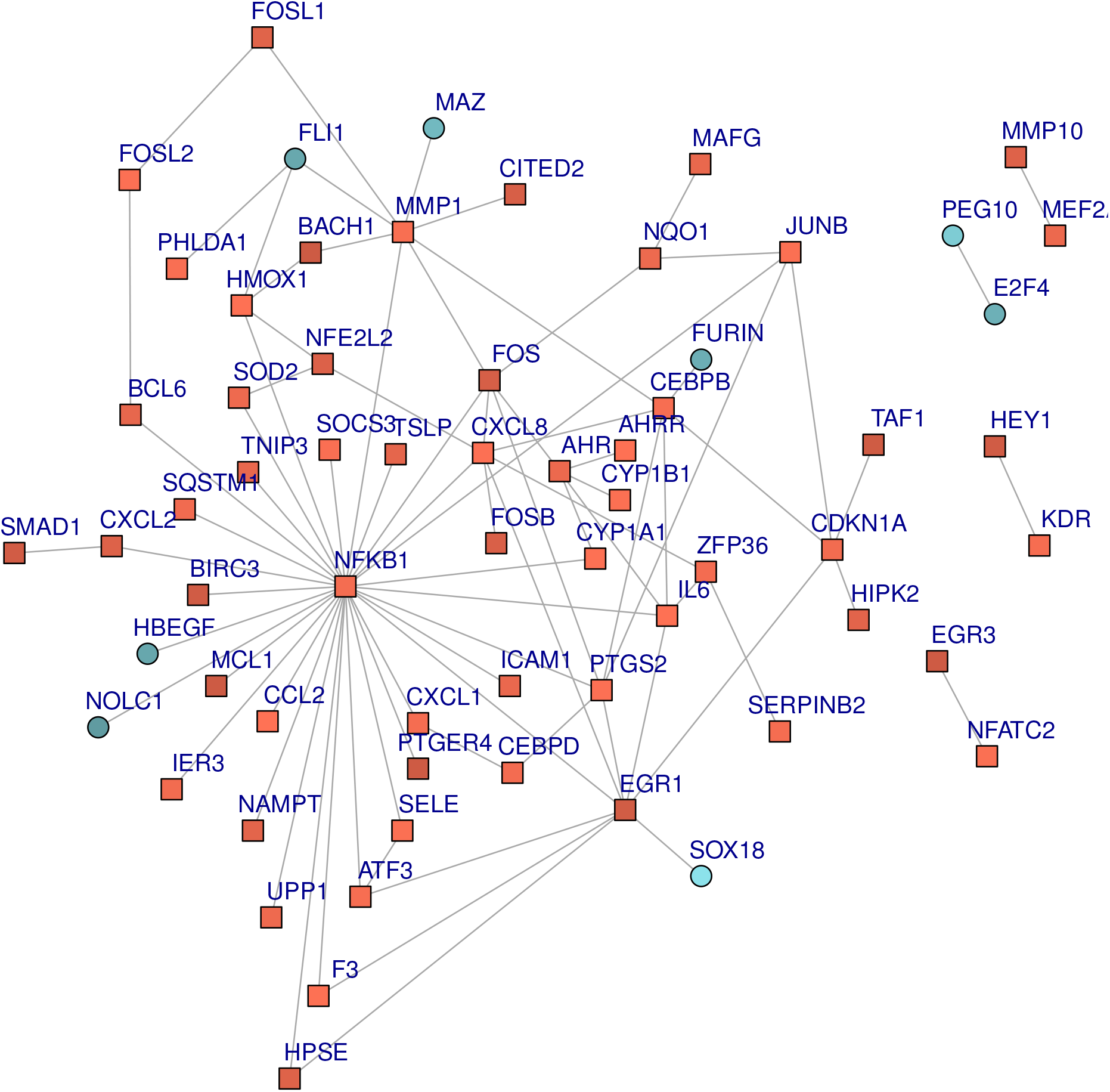
Coherent subnetwork in ultrafine particle-exposure study (FDR α = 0.05). The GRN subnetwork of the human pulmonary endothelial cells undergoing coherent differential expression from exposure to ultrafine particles. Square nodes indicate increased gene expression, while circular nodes indicate decrease in gene expression after exposure. The brighter the nodes, the more significant the change.

## 4 Conclusion

We have shown that our method infers differentially actived modules in gene regulatory networks. While this work is developed in the context of gene regulatory networks, the methodology is sufficiently general for application to other molecular networks. We proposed three procedures for FDR control in order to control the boundary links that do not form part of the DE networks in order to separate active modules. Controlling both genes and links was not previously performed. While the simultaneous procedure needs no further theoretical justification, the two-stage procedures pose a theoretical problem, due to the conditional nature of the second stage. We have presented here simulations demonstrating the non-inferiority of the forward procedure against the simultaneous procedure as empirical validation. We have further demonstrated the utility of the reverse procedure, which is more sensitive than the forward procedure to signal.

Accounting for uncertainty improves inference of differentially activated modules. We have demonstrated the difference between quantile-filtration and error-control in Figure 4. Crucially, quantile-filtration is independent of the proportion of signals, returning similar results for noise and signal scenarios, whereas controlling error implicitly adjusts to the proportion of truly DE genes. With inspiration from this previous work, we reframed the differential link score in a manner permitting full error control. The DLS originates in Warsow *et al*. (2010), where the authors address uncertainty by normalizing the log-fold change in gene expression by Welch’s estimate of the common variance, 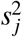, for log-expression each gene, log(*x_ij_*) → log(*x_ij_*)/*s_j_*. Substituting this into (1), the DLS can be interpreted as the sum of two correlated *t*-statistics, effectively amplifying the signal of the correlated components over the uncorrelated components. In a later work, an effort at error control is made by normalizing the DLS using an estimate of the variance 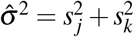 (Warsow *et al*. (2013)). However, this estimate does not account for the covariance between genes *j* and *k*, distorting the significance of the findings. This approach was subsequently abandoned in Ernst *et al*. (2017) for a strictly quantile-based analysis.

None of the four methods F/P, R/P, BioNet, or limma uniformly dominates any other. There are some scenarios where BioNet outperforms the rest; however, BioNet’s extreme variation in sensitivity is hard to predict. Remarking that all methods return a set of genes with increased proportion of signal genes, procedures R/P or BioNet may be appropriate for narrowing the search space while discarding as few signal genes as possible. When it is important that error be well-controlled, the F/P and limma procedures are most appropriate. We recommend comparing network-constrained approaches to a limma baseline. The disparity between these can indicate whether we are observing a scattered or modular differential expression regime. Further, it can safeguard against high variation in precision (R/P and BioNet) or high variation in sensitivity (BioNet).

Network-constrained approaches to identifying regulated features are accompanied with the desire for increased power to detect the truly relevant changes before an expansive, noisy background. However, the network-constraints at the same time bias our possible results. It may be that the non-coherent differential signals in the data point to interesting regulatory changes. These could represent cases where the actual regulatory structure has changed. Thus, it is worthwhile to investigate both coherent and non-coherent differential regulatory signals.

## Acknowledgements

The authors would like to thank Dr Georg Fuellen and Frida Gorreja for critical review of this article.

